# Acyloxyacyl Hydrolase Modulates the Gut Microbiome Through Transcriptional Regulators of Corticotropin-Releasing Factor

**DOI:** 10.1101/2021.03.04.433950

**Authors:** Afrida Rahman-Enyart, Lizath M. Aguiniga, Wenbin Yang, Ryan E. Yaggie, Bryan White, Michael Welge, Loretta Auvil, Matthew Berry, Colleen Bushell, Anthony J. Schaeffer, David J. Klumpp

## Abstract

Gut microbiome-host interactions play a crucial role in health and disease. Altered gut microbiome composition has been observed in patients with interstitial cystitis/bladder pain syndrome (IC/BPS), a disorder characterized by pelvic pain, voiding dysfunction, and often co-morbid with anxiety/depression. We recently showed that mice deficient for acyloxyacyl hydrolase (AOAH) mimic pelvic pain symptoms and comorbidities of IC/BPS and also exhibit gut dysbiosis. In addition, we previously identified that the conditional knockout (cKO) of two transcriptional regulators of the gene encoding corticotropin-releasing factor, *Crf*, that are downstream of AOAH, aryl hydrocarbon receptor (AhR) and peroxisome proliferator-activated receptor-*γ* (PPAR_*γ*_), alleviate anxiety/depressive and voiding phenotypes of AOAH-deficient mice. Here, we examined the effects of AhR and PPAR_*γ*_ in CRF-expressing cells on the dysbiosis of AOAH-deficiency. AOAH-deficient mice with cKO of *PPAR_γ_* and *AhR/PPAR_γ_* exhibited reduced pelvic allodynia compared to AOAH-deficient mice, suggesting a role for PPAR_*γ*_ in regulating pelvic pain. 16S rRNA sequencing of fecal stool from female AOAH-deficient mice with a cKO of AhR and/or PPAR_*γ*_ in CRF-expressing cells identified altered gut microbiota distinct from AOAH-deficient stool. The cKO of AhR and PPAR_*γ*_ showed improved cecum barrier function in females compared to AOAH-deficient mice, whereas males were primarily affected by PPAR_*γ*_, suggesting sex differences in gut responses. Pair-wise comparison of microbiota also suggested sex differences in response to AOAH-deficiency and conditional knockout of *AhR* and *PPAR_γ_*. Our findings suggest that the dysbiosis and leaky gut of AOAH deficiency is mediated by AhR and PPAR_*γ*_ in CRF-expressing cells and reveal a novel mechanism and therapeutic targets for pelvic pain.

## INTRODUCTION

The gut microbiome plays an important role in health and metabolism, and dysbiosis has been associated with numerous diseases and visceral pain (7, 25, 30). It has previously been shown that patients suffering from interstitial cystitis/bladder pain syndrome (IC/BPS or “IC”), a chronic condition characterized by pelvic pain and urinary dysfunction, have altered fecal microbiota (3, 4, 14, 21). IC/BPS continues to be a clinical challenge due to its unknown etiology and lack of effective biomarkers (21). With this goal in mind, we previously conducted a genetic screen to identify modulators of pelvic pain severity and identified a locus encoding *acyloxyacyl hydrolase* (*Aoah*) as a candidate gene (37).

AOAH is a neutrophil lipase best known for its role in the detoxification of bacterial lipopolysaccharides (LPS) through the removal of secondary acyl chains from the lipid A moiety, resulting in the downregulation of host response to bacterial infection and the attenuation of the inflammatory response (10, 12, 18, 22, 31). We have previously demonstrated that AOAH-deficient mice develop a phenotype of spontaneous pelvic pain and are more susceptible to induced pelvic pain models (37). In addition, AOAH-deficient mice mimic key aspects of IC/BPS, including gut dysbiosis and several associated comorbidities (1, 2, 26, 37). Altering the gut microbiome of AOAH-deficient mice by co-housing or gavage of healthy stool slurry alleviates pelvic pain, suggesting a role for gut flora in modulating the pelvic pain phenotype (26).

Corticotropin-releasing factor (CRF) is a well-known regulator of the stress response through the regulation of the hypothalamic-pituitary-adrenal (HPA) axis and is also a key modulator in pain responses. IC/BPS patients exhibit altered diurnal cortisol expression implicating HPA axis dysregulation (19). Previous studies in our lab demonstrated that AOAH-deficient mice exhibit increased expression of central nervous system (CNS) arachidonic acid, increased *Crf* expression in the paraventricular nucleus, and elevated serum corticosterone, consistent with dysregulation of the HPA axis (2). In addition, we identified two arachidonic acid-dependent transcription regulators of *Crf* that play a role in AOAH-deficient phenotypes, aryl hydrocarbon receptor (AhR) and peroxisome proliferator-activated receptor-γ (PPAR_*γ*_) (2). Conditional knockout (cKO) of *AhR* and/or PPAR_*γ*_ in CRF-expressing cells in AOAH-deficient mice rescued phenotypes often observed in IC/BPS patients, such as depressive behavior and voiding dysfunction (1, 2).

CRF also plays a role in regulating the complex communication pathways of the microbiota-gut-brain axis and alterations of CRF expression have been shown to promote changes in gut microbiota composition (16). Since we have observed gut dysbiosis in AOAH-deficient mice and a role for CRF in AOAH-deficient phenotypes, we hypothesized that the cKO of transcriptional regulators of *Crf* would alter gut dysbiosis mediated by AOAH deficiency. We observed that the cKO of *AhR* and/or PPAR_*γ*_ in CRF-expressing cells in AOAH-deficient mice altered the gut microbiome and “leaky gut” phenotype in a sex-dependent manner and cKO of both *AhR* and PPAR_*γ*_ resulted in alleviation of pelvic allodynia in female mice. Our findings suggest that dysbiosis associated with AOAH deficiency is mediated by AhR and PPAR_*γ*_ in CRF-expressing cells and reveal novel therapeutic targets for treating pelvic pain.

## MATERIALS AND METHODS

### Animals

Ten to twelve-week-old male and female wild type (WT) C57BL/6 mice were purchased from The Jackson Laboratory. *Aoah−/−* mice (B6.129S6-Aoahtm1Rsm/J) were a gift from Dr. Robert Munford of NIAID and maintained on a 12h:12h light:dark cycle as previously described (2). Conditional knockout mice were generated as previously described in Aguiniga et al. (2)

### Pelvic allodynia

Pelvic allodynia was measured by measuring responses to von Frey filament stimulation to the pelvic region, as previously described in Rudick et al. (28). Briefly, pelvic allodynia was measured for WT and AOAH-deficient mice with or without conditional knockout of *AhR* and/or *PPAR_γ_*. Mice were placed and allowed to acclimate to the test chamber for 5 min. Starting with the filament that applied the lowest force, five von Frey filaments were applied 10 times to the pelvic region. A behavioral response was measured as painful if the animal jumped, shook the hind paws, or excessively licked the pelvic region.

### Gut microbiome analyses

Fecal gut microbiota were analyzed as previously reported in Rahman-Enyart et al. (26). Briefly, DNA was extracted from fecal pellets from male and female mice using the QIAamp DNA Stool Mini Kit (QIAGEN, Hilden, Germany) and homogenized using 0.1 mm zirconia/silica beads (Biospec, Bartlesville, OK) in 1.4 mL ASL buffer (QIAGEN). Amplicon sequencing of the 16S rRNA V3–V5 hypervariable region was carried out to identify phylotype profiles of microbiota. The V3–V5 hypervariable region was amplified through 30 cycles with primers 357F (CCTACGGGAGGCAGCAG) and 926R (CCGTCAATTCMTTTRAGT). Amplicon pools were quantified on a Qubit fluorimeter, and fragment sizes were determined by Agilent bioanalyzer High Sensitivity DNA LabChip (Agilent Technologies, Wilmington, DE). Amplicons were spiked with a PhiX control library to 20% and mixtures were then sequenced on an Illumina MiSeq V2 (250nt from each end). Sequence reads were binned at 97% identity using QIIME and Galaxy in order to define abundance of OTUs and their phylogenic relationships.

### Transepithelial electrical resistance

Cecum permeability was measured as transepithelial electrical resistance (TEER) as previously described in Rahman-Enyart et al. (26). Ceca were bisected and placed in Ringer’s solution at room temperature followed by mounting onto cassettes with 0.126cm^2^ aperture. Cassettes were inserted into an Ussing chamber (Physiologic Instruments EM-CSYS-2) with KCl saturated salt-bridge electrodes. The chambers were then filled with Ringer’s solution and bubbled with carbogen (95% O_2_/5% CO_2_). The chambers were maintained at 37 °C until resistance was stabilized, about 1 hr. Voltage was clamped and current was passed every 3 s at 30 s intervals using a VCC MC2 multichannel voltage-current clamp amplifier using Acquire and Analyze software v2.3.4 (Physiologic Instruments).

### Statistical analysis

Results were statistically analyzed by implementing either Student’s t-test or one-way analysis of variance (ANOVA) followed by Tukey’s Multiple comparisons test. All statistics were done using Prism software, version 6 (GraphPad, Inc). Differences were considered statistically significant at P<0.05.

## RESULTS

### Conditional knockout of *PPAR_γ_* and *AhR* alleviates pelvic pain in AOAH-deficient mice

AOAH-deficient mice exhibit increased pelvic pain behaviors compared to wild-type controls (37). To determine whether the cKO of *Ahr* or *PPAR_γ_* in CRF-expressing cells could alter pelvic allodynia in female AOAH-deficient mice, we quantified allodynia in response to von Frey filaments applied to the pelvic region (Fig. 1). Similar to our previous findings (37), we observed that AOAH-deficient mice exhibited a robust pelvic pain phenotype. We observed a significant reduction (~40%) in pelvic allodynia in AOAH-deficient mice with a cKO of both *AhR* and *PPAR_γ_*. The cKO of *AhR* only did not alleviate the pelvic pain phenotype in AOAH-deficient mice. We observed a slight reduction in pelvic allodynia (~20%) in AOAH-deficient mice with a cKO of *PPAR_γ_*; however, these results did not approach significance. These data suggest that both AhR and PPAR_*γ*_ are required for the pelvic pain phenotype observed in AOAH-deficient mice and the cKO of just one of these genes is not sufficient to alleviate pelvic allodynia.

**Figure 1.**
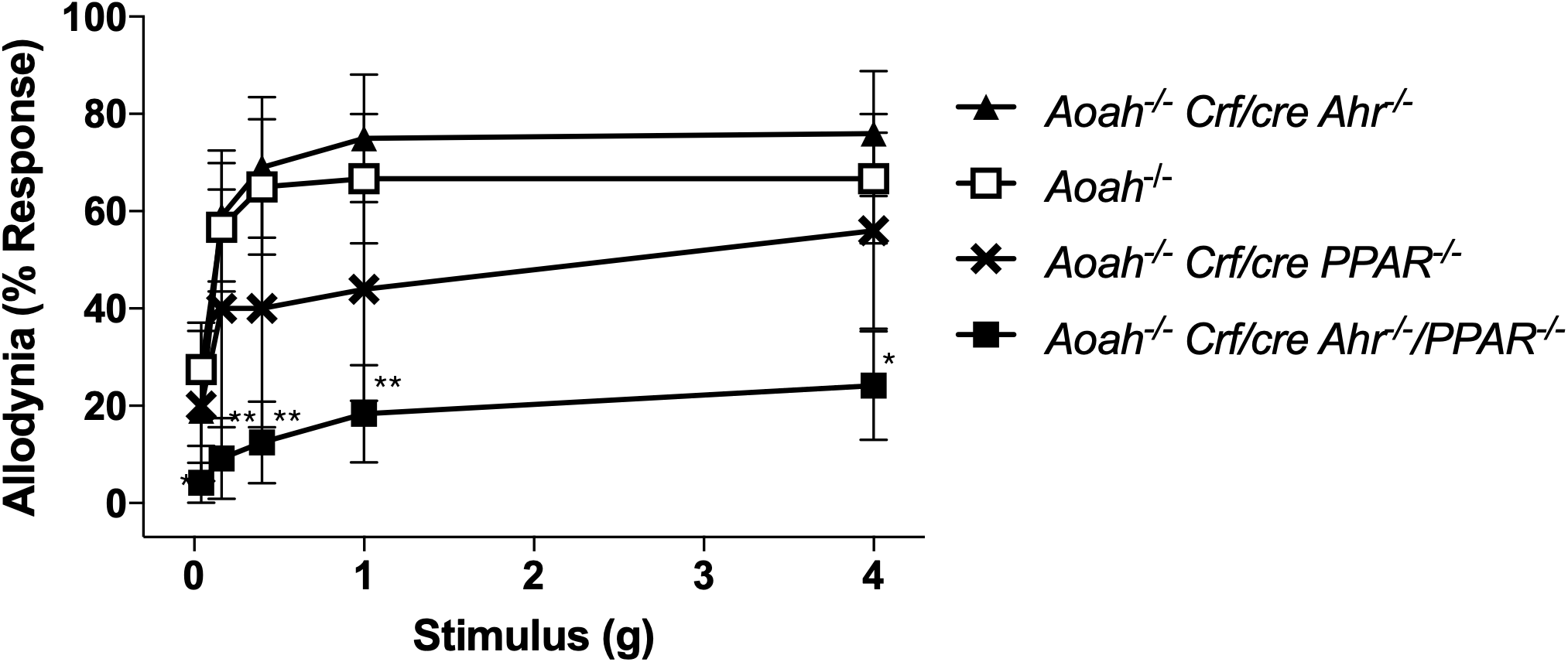
Conditional knockout of *AhR/PPAR_γ_* in female AOAH-deficient mice alleviates pelvic allodynia. AOAH-deficient mice exhibited increased pelvic allodynia compared to AOAH-deficient mice with cKO of *AhR/PPAR_γ_* in CRF-expressing cells, as shown through response to von Frey filaments stimulating the pelvic region (n=5-12 mice; *P<0.05, **P<0.01, Student’s t test, two tailed). Data represented as average response (%) ± SEM.

### Conditional knockout of *PPAR_γ_* and *AhR* alters fecal microbiota

We have previously shown that AOAH-deficient mice exhibit an altered gut microbiome and manipulation of the microbiota alters the pelvic pain phenotype (26). Therefore, we next sought to identify whether the cKO of *Ahr* or *PPAR_γ_* in AOAH-deficient mice would result in a transformed gut microbiome. We utilized 16S rRNA amplicon sequencing data to identify large compositional differences in fecal microbiota between wild type and AOAH-deficient mice with or without cKO of *Ahr* or *PPAR_γ_* (Fig. 2). As shown by principal component analysis (PCA, Fig. 2A) and the corresponding heat map (Fig. 2B), AOAH-deficient mice exhibit an altered gut microbiota compared to wild type control, similar to our previous findings (26). Mice with cKO for *Ahr* or *PPAR_γ_* show group variance along the PC1 (vertical) axis in bacterial composition of gut flora compared to AOAH-deficient and wild type mice, suggesting an altered but unique microbiome (Fig. 2A). We observed that the gut microbiome of AOAH-deficient mice with cKO for *PPAR_γ_* was most similar to wild type control, whereas cKO of *AhR* was more similar to AOAH-deficient microbiota. The cKO of *PPAR_γ_* in wild type mice also showed an altered microbiome, suggesting a role for *PPAR_γ_* in regulating bacterial composition of gut flora. Overall, these data identify a role for PPAR_*γ*_ in regulating the gut dysbiosis phenotype in AOAH-deficient mice.

**Figure 2.**
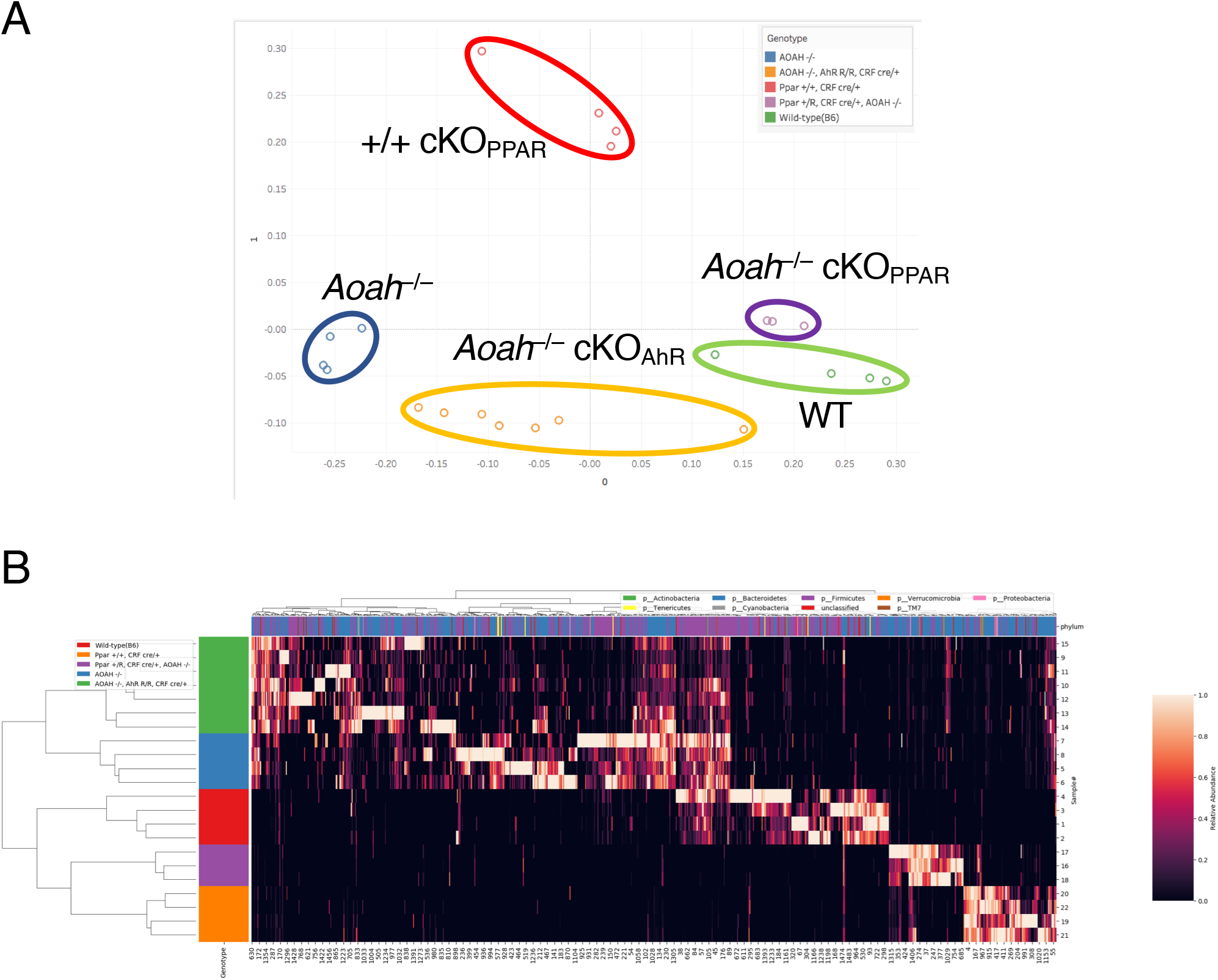
Conditional knockout of *AhR* or *PPAR_γ_* in female AOAH-deficient mice alters gut microbiome composition. **A**. PCA plot of 16S rRNA analyses of fecal stool in female wild-type and AOAH-deficient mice and mice with a cKO of *AhR* or *PPAR_γ_* in CRF-expressing cells. Dots represent individual mice; AOAH-deficient mice shown in blue, wild-type mice shown in green, AOAH-deficient mice with cKO of *Ahr* shown in yellow, AOAH-deficient mice with cKO of *PPAR_γ_* shown in purple, WT mice with cKO of *PPAR_γ_* shown in red (n=3-6 mice). **B**. Heat map of 16S rRNA analyses of fecal stool from AOAH-deficient (blue), wild-type (red), AOAH-deficient/cKO of *Ahr* (green), AOAH-deficient/cKO of *PPAR_γ_* (purple), and WT/cKO of *PPAR_γ_* (orange) female mice (n=3-6 mice)

### Conditional knockout of *PPAR_γ_* and *AhR* alters gut permeability in AOAH-deficient mice

Previous studies have linked gut dysbiosis and gut permeability (9, 20, 36), and we have observed a “leaky gut” phenotype in AOAH-deficient mice (26). To identify whether AhR and PPAR_*γ*_ play a role in gut “leakiness,” we assessed cecum barrier function in male and female wild type and AOAH-deficient mice with cKO of *Ahr* and/or *PPAR_γ_* by quantifying cecum transepithelial electrical resistance (TEER) ex vivo via an Üssing chamber (Fig. 3A and B, respectively). As we have previously reported (26), cecum TEER was significantly lower in AOAH-deficient mice compared to wild type in both males (27.66 ± 6.63 Ω•cm^2^ for wild type and 17.11 ± 6.11 Ω•cm^2^ for *Aoah−/−*; Fig. 3A) and females (33.58 ± 4.77 Ω•cm^2^ for wild type and 19.94 ± 6.66 Ω•cm^2^ for *Aoah−/−*; Fig. 3B). The cKO of *PPAR_γ_* or both *AhR/PPAR_γ_* in male AOAH-deficient mice resulted in a significant increase in TEER (36.25 ± 4.12 Ω•cm^2^ for *PPAR_γ_* cKO and 35.10 ± 4.36 Ω•cm^2^ for *Ahr*/*PPAR_γ_* cKO; Fig. 3A), compared to male AOAH-deficient mice. In contrast, in female mice, the cKO of *AhR* or both *AhR/PPAR_γ_* resulted in a significant increase in TEER (29.90 ± 3.79 Ω•cm^2^ for *AhR* cKO and 29.03 ± 6.31 Ω•cm^2^ for *Ahr*/*PPAR_γ_* cKO; Fig. 3B), compared to female AOAH-deficient mice. These data exhibit that both AhR and PPAR_*γ*_ play a role in regulating gut permeability in AOAH-deficient mice, and do so in a sex-dependent manner.

**Figure 3.**
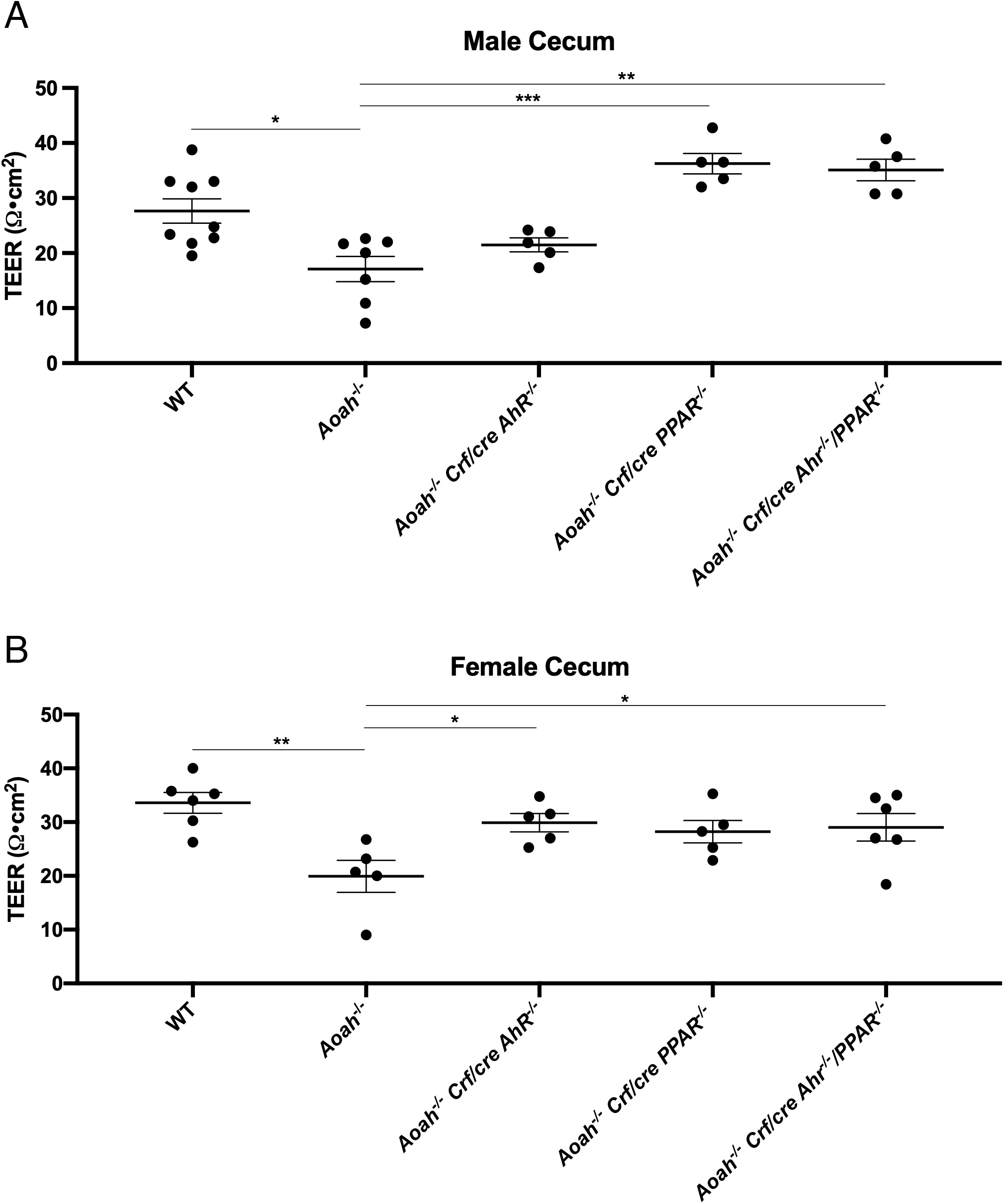
Conditional knockout of *AhR* or *PPAR_γ_* in AOAH-deficient mice alters TEER. Ceca of male **(A)** and female **(B)** mice were measured for transepithelial electrical resistance (TEER). Both male and female AOAH-deficient mice exhibited lower TEER compared to WT. **A**. The cKO of *PPAR_γ_* or *Ahr*/*PPAR_γ_* in male AOAH-deficient mice showed improved TEER compared to AOAH-deficient mice. **B**. The cKO of *Ahr* or *Ahr*/*PPAR_γ_* in female AOAH-deficient mice showed improved TEER compared to AOAH-deficient mice. (n=5-9 mice; *P<0.05, **P<0.01, ***P<0.001, One-Way ANOVA followed by post-hoc Tukey HSD). Data represented as individual values (dots) and average (represented by horizontal line) ± SEM

### *PPAR_γ_* and *AhR* regulate sex-dependent alterations in AOAH-deficient microbiota

Since our observations indicate sex differences in gut permeability regulated by AhR and PPAR_*γ*_ (Fig. 3), we next addressed whether bacterial composition of the gut would be altered in a sex-dependent manner after cKO of *AhR* and/or *PPAR_γ_* in AOAH-deficient mice. Corroborating our data from Fig. 2, 16S rRNA amplicon sequencing indicated that the cKO of *AhR* or *PPAR_γ_* in female AOAH-deficient mice resulted in a gut microbiome that was distinct from female AOAH-deficient mice (Fig. 4, left column). In addition, we also observed that the cKO of both *Ahr*/*PPAR_γ_* resulted in an altered fecal microbiota in female AOAH-deficient mice. Female AOAH-deficient mice with cKO for *PPAR_γ_* or both *Ahr*/*PPAR_γ_* show a larger group variance along the PC1 (vertical) axis in bacterial composition of gut flora compared to female AOAH-deficient mice and AOAH-deficient mice with cKO for only *AhR*. These data suggest that both AhR and, to a greater degree, PPAR_*γ*_ play a role in regulating the gut microbiome in female AOAH-deficient mice.

**Figure 4.**
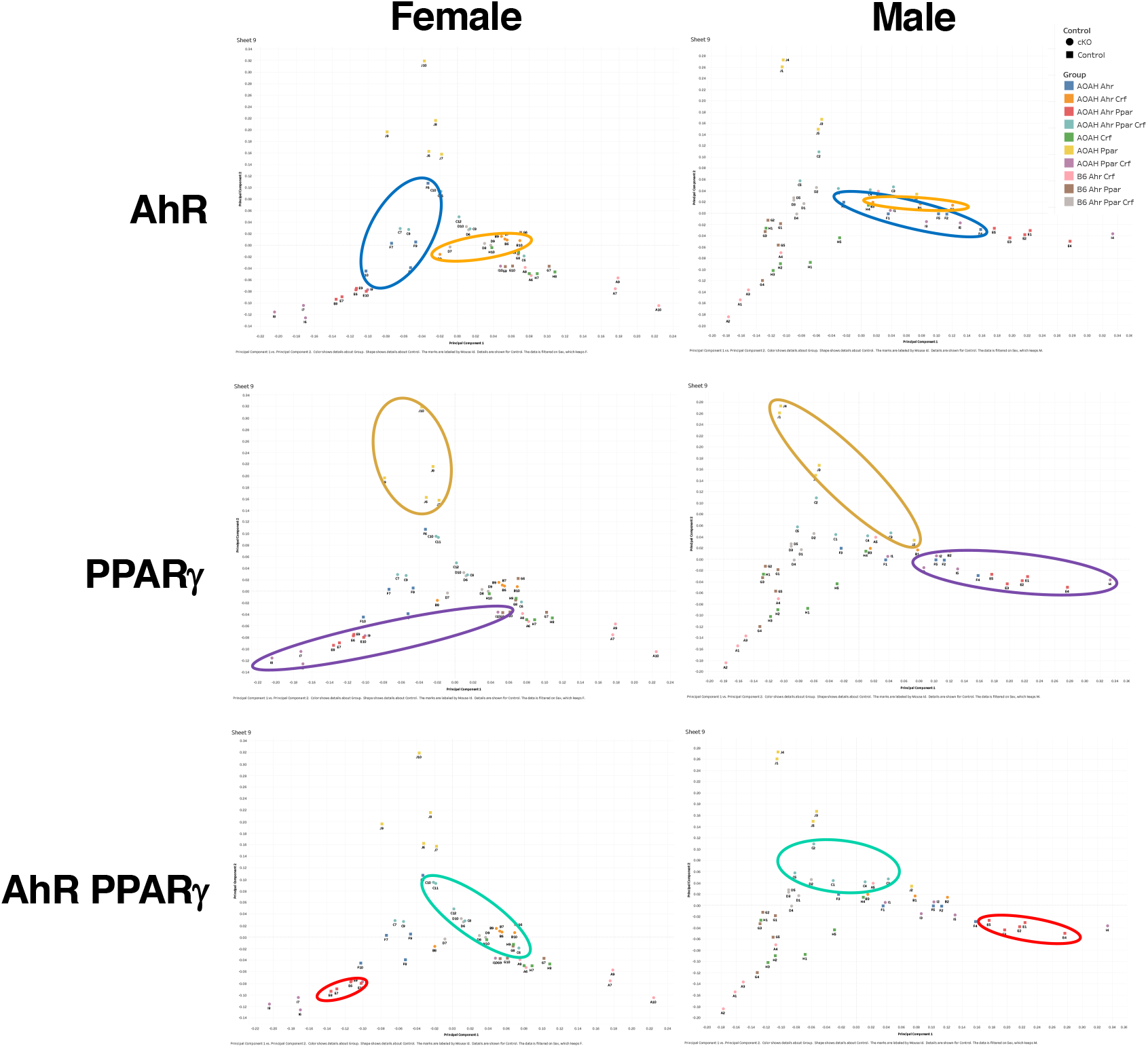
Sex differences in microbiome composition in AOAH-deficient mice with conditional knockout of *AhR* or *PPAR_γ_*. PCA plots of 16S rRNA analyses of fecal stool in female (left column) or male (right column) AOAH-deficient mice with a cKO of *AhR* (top), *PPAR_γ_* (middle), or *AhR*/*PPAR_γ_* (bottom). Dots represent individual mice; *Aoah−/− Ahr* control shown in blue and cKO shown in orange, *Aoah−/− PPAR_γ_* control shown in yellow and cKO shown in purple, *Aoah−/− Ahr/PPAR_γ_* control shown in red and cKO shown in green (n=5 mice for all conditions)

We next utilized 16S rRNA amplicon sequencing in male AOAH-deficient mice to identify whether the conditional knockout of *AhR* and/or *PPAR_γ_* would result in altered gut bacterial composition similar to our observations in females. In contrast to female mice, we observed that in male AOAH-deficient mice the cKO of *PPAR_γ_* or both *Ahr*/*PPAR_γ_*, but not *AhR* only, resulted in a gut microbiome that was distinct from AOAH-deficient mice (Fig. 4, right column). These findings suggest a sex difference in the role of AhR and PPAR_*γ*_ in regulating gut bacterial composition, where both AhR and PPAR_*γ*_ play a role in female gut flora but only PPAR_*γ*_ play a role in male flora.

To determine the specificity of AhR and PPAR_*γ*_ modulation of AOAH-deficient gut microbiota, we examined the relative abundance of bacterial phyla (Fig. 5). As expected, the majority of bacteria identified in all mouse groups belonged to Bacteroidetes and Firmicutes phyla. Similar to our previous findings (26), we did not observe differences in mean bacterial abundance belonging to Bacteroidetes and Firmicutes phyla in wild type and AOAH-deficient conditions, but did observe differences within less abundant phyla (Fig. 5). In contrast, we observed that the cKO of *PPAR_γ_* or both *Ahr*/*PPAR_γ_* resulted in a decrease in Firmicutes phyla when compared to AOAH-deficient and wild type mice (Fig. 5 C-F). We observed an increase in abundance in Bacteroidetes phyla in male AOAH-deficient mice with cKO of *Ahr*/*PPAR_γ_,* but not in females (Fig. 5E and F). In addition, the cKO of *AhR* resulted in greater abundance of the less common Cyanobacteria and Proteobacteria phyla in both males and females when compared to AOAH-deficient and wild type mice (Fig. 5A and B). We observed sex differences in the abundance of bacteria belonging to Actinobacteria phyla, where male AOAH-deficient mice with cKO for *Ahr* and/or *PPAR_γ_* exhibited greater abundance compared to AOAH-deficient and wild type mice, which was not observed in females (Fig. 5B, D, and F compared to Fig. 5A, C, and E).

**Figure 5.**
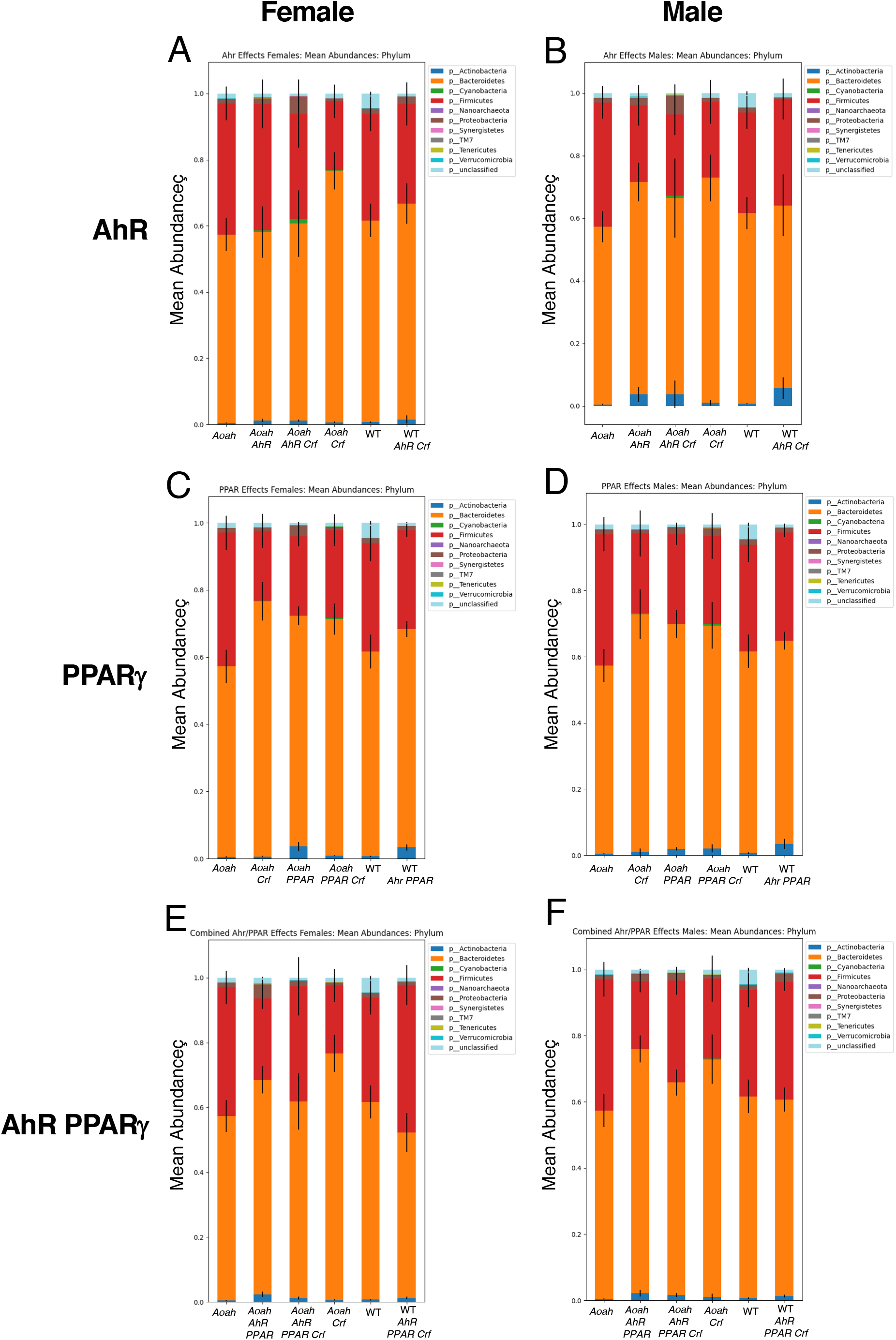
Quantification of mean abundance of bacterial phyla in female **(A, C, and E)** and male **(B, D, and F)** wild-type, AOAH-deficient, and cKO mice using 16S rRNA sequencing (n=5 mice for all conditions)

In addition, we analyzed relative abundance of Firmicutes and Bacteroidetes phyla in response to cKO of *AhR*, *PPAR_γ_*, or both *Ahr*/*PPAR_γ_.* Relative abundance of bacteria belonging to Firmicutes phyla was greater in AOAH-deficient mice compared to WT mice; whereas, bacteria belonging to Bacteroidetes phyla was more abundant in WT compared to AOAH-deficient mice (Fig. 6). We observed that the cKO of *AhR*, *PPAR_γ_*, or both *Ahr*/*PPAR_γ_* altered relative abundance of both Firmicutes and Bacteroidetes phyla to abundance that more closely resembled wild type. Of note, in females, we observed a greater change in mice with a conditional knockout for *PPAR_γ_* compared to *AhR* (Fig. 6E and F compared to Fig.6A and B). Overall, these data show that both AhR and PPAR_*γ*_ play important roles in regulating the gut microbiome in AOAH-deficient mice, and their effects are sex-dependent.

**Figure 6.**
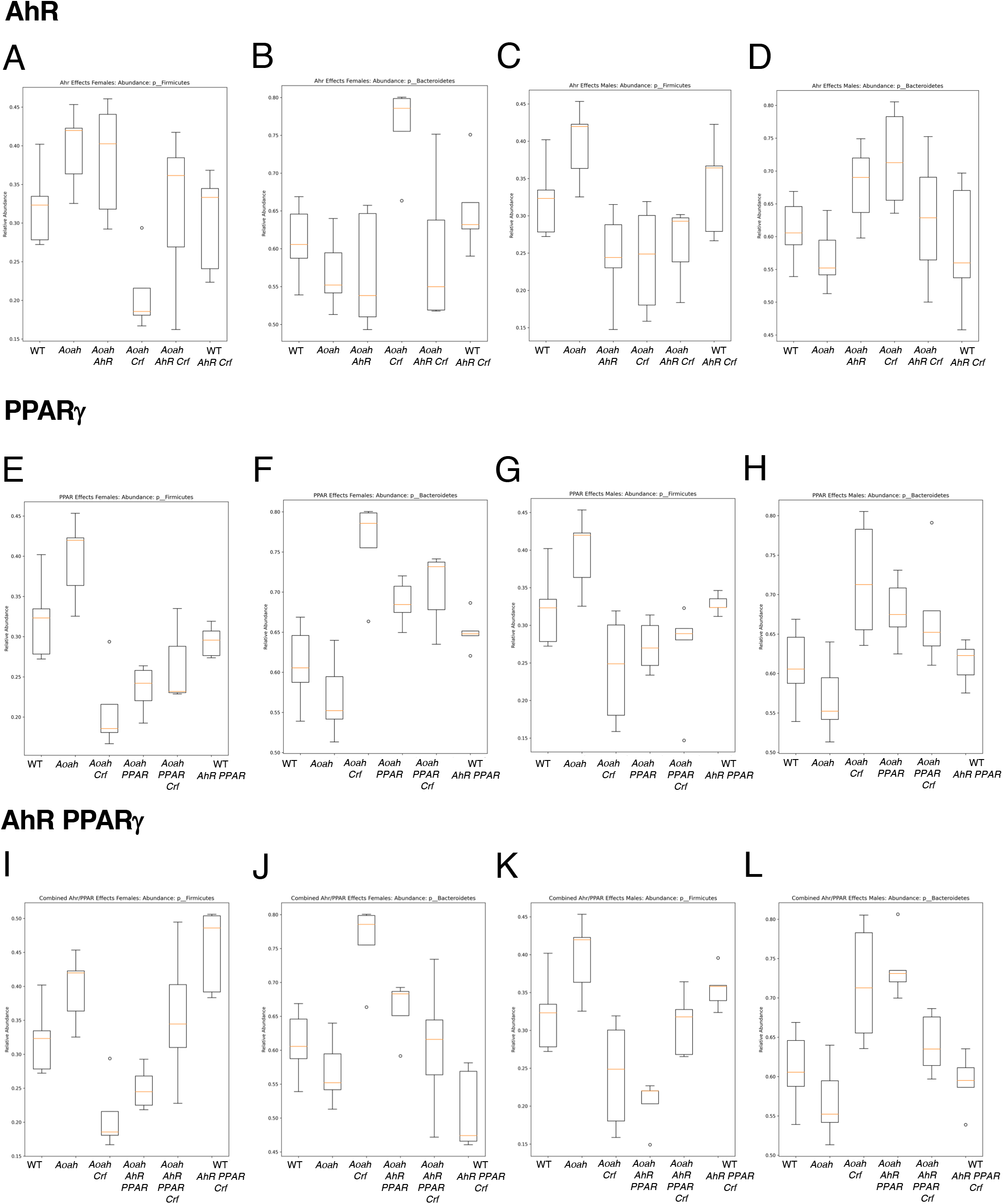
Relative abundance of Bacteroidetes and Firmicutes phyla in AOAH-deficient stool with conditional knockout of *AhR* and/or *PPAR_γ_*. The effects on cKO of *AhR* **(A-D)**, *PPAR_γ_* **(E-H)**, or *AhR/PPAR_γ_* **(I-L)** on relative abundance of bacteria belonging to Bacteroidetes and Firmicutes phyla was measured in WT and AOAH-deficient female **(A, B, E, F, I, and J)** and male **(C, D, G, H, K and L)** mice using 16S rRNA sequencing (n=5 for all conditions)

## DISCUSSION

AOAH deficiency recapitulates several aspects of IC/BPS and we have previously identified *Aoah* as a genetic modulator of gut microbiota composition (1, 2, 26, 37). Here we show that AhR and PPAR_*γ*_ in CRF-expressing cells can modulate gut dysbiosis mediated by AOAH deficiency. CRF is best known for its role in the stress response through regulating the HPA axis (24). In addition, CRF also plays a crucial role in pain modulation, gut function, and composition of the gut microbiome (11, 16, 33, 34). We have previously shown increased *Crf* expression in the paraventricular nucleus of AOAH-deficient mice as well as elevated serum corticosterone (2). In addition, cKO of *Ahr* and *PPAR_γ_*, two lipid-responsive transcription regulators of *Crf*, can alleviate voiding dysfunction and depressive behaviors in AOAH-deficient mice (1, 2). Increased activation of mast cell CRFR1 receptors via elevated CRF in the enteric epithelium has been observed to increase epithelial permeability (27), a phenotype often observed in gut dysbiosis (13). Consistent with these data, we observed increased leakiness in ceca of AOAH-deficient mice that improved upon the cKO of *AhR* or *PPAR_γ_* (Fig. 3), suggesting a role for CRF in gut epithelial permeability in AOAH-deficient mice.

Although we observed changes in pelvic allodynia, gut microbiome composition, and TEER upon manipulation of *Crf* expression in AOAH-deficient mice, we identified that the effects of *AhR, PPAR_γ_*, or both *AhR/PPAR_γ_* was dependent on phenotype and sex. The cKO of *PPAR_γ_* had a greater role in improving TEER and altering the gut microbiome in males compared to the cKO of *AhR* (Figs. 3 and 4). In female AOAH-deficient mice, both AhR and PPAR_*γ*_ play a role in altering the gut microbiota (Fig. 4), but AhR is the key driver in modulating gut permeability (Fig. 3). Additionally, the cKO of both *AhR/ PPAR_γ_* was required to alleviate the pelvic pain phenotype in females (Fig. 1). Our findings suggest that both AhR and PPAR_*γ*_ are regulators of gut function but possess unique roles that are sex-dependent. Sex-specific differences in gut microbiota have been observed in both mice and humans (15). Unsurprisingly, sex hormones such as testosterone and estrogen are directly linked to altered gut microbiota. In addition, metabolism, body mass index, and colonic transit time differ amongst sex and can also alter gut flora (15). Whether AhR and PPAR_*γ*_-dependent transcription of *Crf* modulates these sex-dependent biological functions in AOAH-deficient mice will require further investigation.

Manipulation of PPAR_*γ*_ and AhR signaling has been associated with microbiota alterations and gut permeability (5, 23). Depletion of butyrate-producing microbes reduces PPAR_*γ*_ signaling in epithelial cells of the colon; whereas, induction of microbiota-dependent PPAR_*γ*_ signaling drives homeostasis by preventing the dysbiotic expansion of Enterobacteriacae (phylum: Proteobacteria) (5). Our findings did not indicate altered Proteobacteria in AOAH-deficient mice with cKO for *PPAR_γ_*; however, we did see increase abundance in Proteobacteria in *AhR* cKOs (Fig. 5), suggesting that AhR is the driving factor for the presence of Proteobacteria in AOAH-deficient mice.

Furthermore, we observed altered mean abundance in bacteria belonging to Firmicutes phylum in AOAH-deficient mice with a cKO of *PPAR_γ_* or both *Ahr/PPAR_γ_* (Fig. 5). A previous study demonstrated that mice fed a high-fat diet showed drastic changes in microbiota composition, including an increase in abundance of Firmicutes as well as dysregulation of the PPAR_*γ*_ pathway (35). In addition, oral administration of water-insoluble polysaccharides to treat a mouse model of alcoholic hepatic steatosis demonstrated both an increase in abundance of Firmicutes and PPAR_*γ*_ signaling (32), suggesting a correlation between expression of Firmicutes and PPAR_*γ*_ signaling. In conjunction to these findings, we observed increased relative abundance of bacteria belonging to Firmicutes phylum in AOAH-deficient mice, which was lower in AOAH-deficient mice with a cKO of *PPAR_γ_* (Fig. 6). Interestingly, acute pain perception has also been associated with altered gut composition, including increase in Firmicutes abundance (29), suggesting a role for bacteria belonging to Firmicutes phylum in pain. Indeed, our findings showed that the cKO of both *Ahr/PPAR_γ_* resulted in a significant decrease in pelvic allodynia in AOAH-deficient mice (Fig. 1).

Metabolites derived from the metabolism of tryptophan have been shown to signal through AhR, and aberrant production of tryptophan-based AhR ligands have been observed in the pathogenesis of inflammatory bowel disease (17, 38). Indeed, we observed that both AhR and PPAR_*γ*_ can alter gut microbiota composition (Figs. 2 and Figs. 4–6). Additionally, we have previously identified the tryptophan metabolite xanthurenic acid as a gut metabolite overexpressed in AOAH-deficient cecal stool (26). Therefore, increased AhR signaling in AOAH-deficient mice through ligands derived from tryptophan metabolism may be driving phenotypes associated with AOAH deficiency. Our current studies did not identify possible alterations in gut metabolites as a result of cKO of *AhR* and *PPAR_γ_* and would be required to assess whether tryptophan metabolism or other gut metabolites may play a role in AhR and PPAR_γ_-mediated gut function and epithelial permeability.

Growing evidence has linked dysbiosis of the gut microbiota with the pathogenesis of several diseases (8). Patients with IC/BPS have altered gut microbiota, a phenotype that AOAH-deficient mice also share (4, 26). As the gut microbiome is involved in bidirectional communication with the CNS, unsurprisingly, several neurological disorders such as depression or neuropathic pain are also affected by an altered gut microbiota (6, 11). Studies from our lab have shown that manipulating the gut microbiome in AOAH-deficient mice improves phenotypes observed in IC/BPS, including pelvic pain and depressive behaviors (26). Here we showed that AOAH-deficient gut dysbiosis and pelvic pain can be manipulated through *Crf* transcription regulators AhR and PPAR_γ_. These findings are central in our understanding of how gut dysbiosis may be modulating the pelvic pain phenotype in IC/BPS. Both AOAH-deficient mice and IC/BPS exhibit HPA dysregulation (2, 19), suggesting a role for CRF in symptomatology. Therefore, future patient studies addressing the correlation of HPA dysregulation and gut dysbiosis will be an interesting avenue for clinical advances in treating IC/BPS.

In summary, the data presented here show that the gut dysbiosis and compromised gut epithelia exhibited by AOAH-deficient mice is modulated by the transcription regulators of *Crf*, AhR and PPAR_γ_. The combined function of AhR and PPAR_*γ*_ also play a role in the pelvic pain phenotype observed in AOAH-deficient mice. These findings demonstrate that the gut microbiome and CRF signaling pathways are promising therapeutic targets for the development of IC/BPS treatments.

## ETHICS APPROVAL AND CONSENT TO PARTICIPATE

All animals were maintained at the Center for Comparative Medicine at Northwestern University and utilized for experimentation under Northwestern IACUC approved protocols.

## AUTHOR CONTRIBUTIONS

L.M.A., W.Y., A.J.S., and D.J.K conceived and designed research; L.M.A., W.Y., R.E.Y. B.W., M.W., L.A., M.B., and C.B. performed experiments; A.R.-E., W.Y., and R.E.Y. analyzed data; A.R.-E., A.J.S., and D.J.K. interpreted results of experiments; A.R.-E., R.E.Y., and D.J.K. prepared figures; A.R.-E. and D.J.K. drafted manuscript; A.R.-E., A.J.S., and D.J.K. edited and revised manuscript; A.R.-E., L.M.A., W.Y., R.E.Y. B.W., M.W., L.A., M.B., C.B., A.J.S., and D.J.K. approved final version of manuscript.

## ACKNOWLEDGEMENTS

We thank Dr. Robert Munford for generously providing AOAH-deficient mice and for many helpful discussions.

